# Astrocyte-derived extracellular matrix proteins regulate synapse remodeling in stress-induced depression

**DOI:** 10.1101/2024.12.30.630684

**Authors:** Ran Zhang, Patricia Rodriguez-Rodriguez, Lucian Medrihan, Jerry C. Chang, Tatiana Ferraro, Pedro Del Cioppo Vasques, Wei Wang, Caroline Menard, Flurin Cathomas, Kenny L. Chan, Lyonna Parise, Shai Shaham, Jean-Pierre Roussarie, Olga Troyanskaya, Scott J. Russo, Ana Milosevic

## Abstract

Major depressive disorder (MDD) is a common mood condition affecting multiple brain regions and cell types. Changes in astrocyte function contribute to depressive-like behaviors. However, while neuronal mechanisms driving MDD have been studied in some detail, molecular mechanisms by which astrocytes promote depression have not been extensively explored. To uncover astrocyte contributions to MDD, we subjected male mice to chronic social defeat stress precipitated by encounters with a dominant male. Animals exposed to this treatment exhibit symptoms indicative of MDD, including reduced social interactions, anxiety, despair, and anhedonia. We then measured astrocyte translating mRNA expression changes in mice that underwent chronic social defeat and control animals using ribosome affinity purification. Bioinformatic analyses reveal significant alterations in the prefrontal cortex (PFC), consistent with previous studies implicating this brain region in MDD. Expression of genes encoding extracellular matrix (ECM) proteins, cell-cell interaction proteins, and proteins controlling glutamatergic synaptic function are significantly altered. These changes correlate with perturbation of glutamatergic transmission, measured by electrophysiology, and increased synaptic cleft size. Among ECM genes, increased expression of mRNA encoding the synaptic remodeling protein **<u>s</u>**ecreted **<u>p</u>**rotein **<u>a</u>**cidic and **<u>r</u>**ich in **<u>c</u>**ysteine (*Sparc*) correlates the most with the depressive phenotype. Furthermore, presence of SPARC and other ECM proteins in synaptosomes is also increased and overexpressing *Sparc* in PFC partially alleviates stress symptoms. Our results raise the possibility that increased expression of *Sparc* may be a natural protective mechanism against stress-induced synaptic dysfunction in depression.

## Introduction

Major depressive disorder (MDD), characterized by anhedonia, helplessness, despair, abnormal food intake, and changes in sleep patterns, and often comorbid with other mood and anxiety disorder, is debilitating to afflicted individuals and has profound societal impact. Although changes on circuitry between various brain regions including prefrontal cortex (PFC), nucleus accumbens (NAc), caudate putamen (CPu), and hippocampus (HIP) have been linked to MDD^1–3^, the contributions of different brain regions and cell types to MDD pathogenesis is still not well understood, while the molecular pathways in specific cell subtypes at play are even less characterized. The neuronal pathophysiology of MDD has been extensively studied, but much less is known about glial contributions to its onset and progression. Strikingly, depression in mice can be induced by ablating astrocytes in the PFC ^4^ or by blockade of astrocyte glutamate uptake ^5^, but not by neuronal ablation ^4^. Thus, astrocytes may play a prominent role in the onset of depressive-like behaviors. Upregulation of glial fibrillary acidic protein (GFAP), a reactive astrogliosis marker, perturbations in the glutamate uptake cycle, and deficient gliotransmission have been reported in MDD patients and mouse models ^6–8^, suggesting causal roles for astrocytes in the progression of the disorder. Consistent with this idea, depressive-like behavior in mice is elicited by astrocyte-specific knockdown of glutamate transporters ^9^, by release of astrocyte-derived ATP in the PFC ^10^, by disruption of astrocyte connexin gap junction ^11,12^, or by perturbation of the astrocyte-specific potassium channel Kir4.1 in the lateral habenula ^13^. Expression of astrocyte-specific genes is altered in post-mortem brain tissue of depression patients ^14,15^, raising the possibility that these changes in astrocyte gene expression may indeed underlie the disease onset.

Similarities in brain architecture, cell types, and genomes, between rodents and humans suggest that mouse models can be used to explore the pathophysiology of MDD and shed light on mechanisms with clinical relevance. In humans, stress can precipitate depressive symptoms and even suicide ^16–20^. Likewise, in rodents, psychological stress activates circuits that lead to depressive behaviors ^21–27^. Indeed, exposure of male mice to a dominant, aggressive male at regular intervals elicits a social defeat-induced stress response that has been used as a model for MDD ^22,28^. In this paradigm, animals exposed to aggressive cage partners develop reduced social interaction, weight loss, and anhedonia ^22,29^.

Here, we used social defeat to explore the role of astrocytes in stress-induced depression in mice. Since astrocytes are heterogeneous in morphology and gene expression and exhibit regional variations in their contribution to synaptic homeostasis both during development and in the adult ^30–37^ we studied multiple brain regions with known role in MDD. These regions were dissected and subjected to translating ribosome affinity purification (TRAP) method to determine gene expression changes between mice that went through chronic social defeat and age-matched controls. Our results uncovered significant changes in the transcriptomes of PFC astrocytes associated with depressive-like behavior. Specifically, we found a highly significant alteration in the expression of genes encoding extracellular matrix (ECM) proteins, especially those supporting glutamatergic synapse function. Among ECM proteins, changes in expression of **<u>s</u>**ecreted **<u>p</u>**rotein **<u>a</u>**cidic and **<u>r</u>**ich in **<u>c</u>**ysteine (*Sparc*), a synaptic ECM protein, were notable. We found that these expression changes correlated with alterations in synapse morphology and function. Importantly, while reducing *Sparc* expression did not rescue the depressive state, over-expression of *Sparc* in the PFC of a Sparc-KO mice alleviated some of the depressive symptoms. Our results, together with previously published findings, ^16,38–44^ suggest that astrocytes may activate protective responses in response to depression-associated synaptic changes.

## Results

### Chronic social defeat alters extracellular matrix and synapse function gene expression in prefrontal cortex astrocytes

To study the involvement of astrocytes in MDD pathophysiology, mice carrying an GFP-L10a sequence under the promotor of Aldh1L1 gene were exposed to a chronic social defeat stress paradigm ^28^. This transgene encodes an astrocyte-specific GFP-tagged ribosomal protein allowing immunoprecipitation of ribosomes and isolation of bound, actively translated mRNAs ^45^. Two days following the last day of stress exposure, animals were assessed by social interaction behavioral test, and a social interaction index (SI) was computed by measuring time spent interacting with a novel mouse (Fig. 1A, B). Mice exposed to chronic stress were either unaffected (resistant to stress, RS, SI>1 similar to control (CTR) mice) or exhibited stress-dependent social avoidance (stress susceptible, SS, SI <1) (Fig. 1A). Immediately following behavior assessment, CTR, SS, and RS mice were sacrificed, and astrocyte ribosome-bound mRNA was isolated from dissected PFC, NAc, CPu, and HIP.

**Figure 1.**
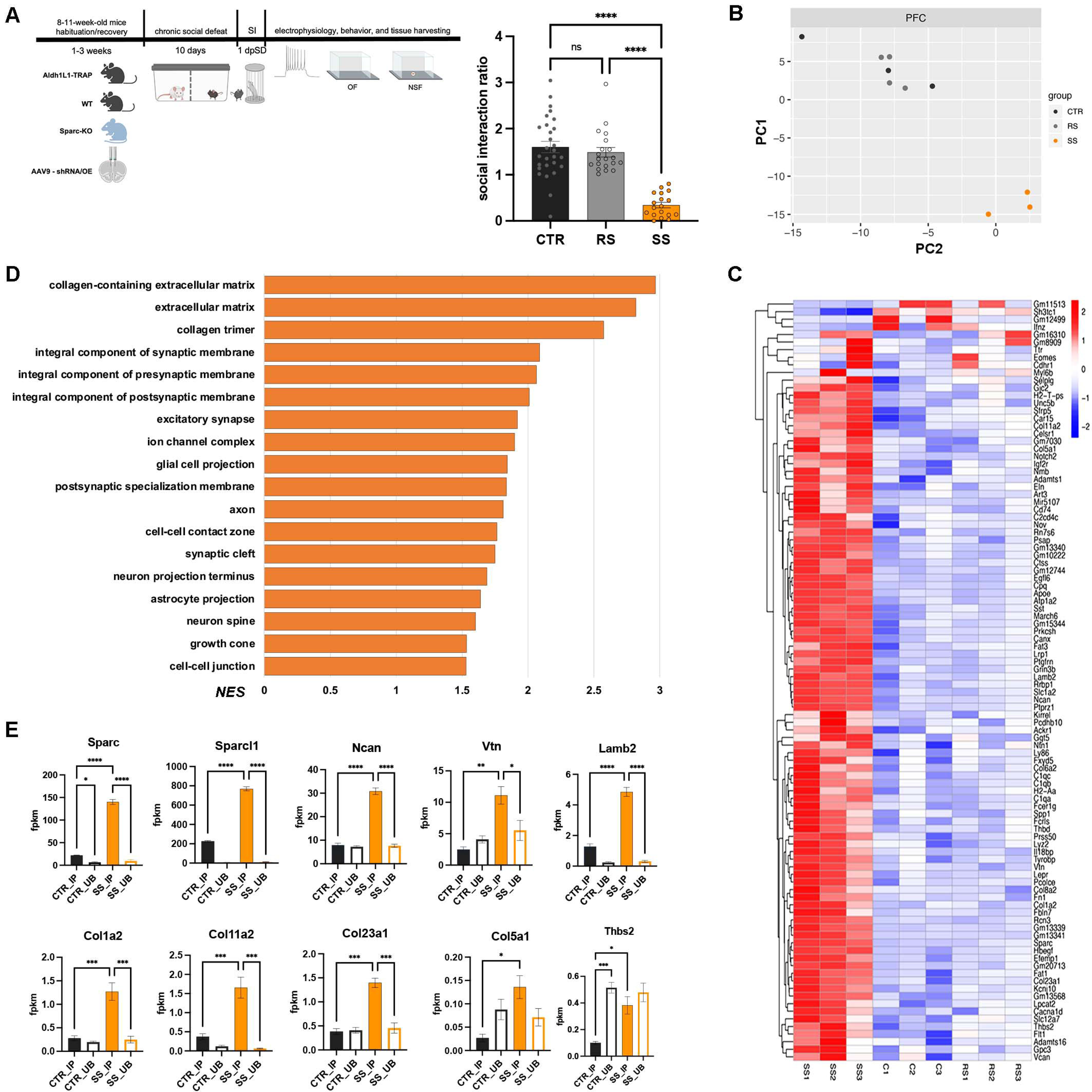
Astrocytes increase expression of extracellular scaffold genes in stress-susceptible mice after chronic social defeat, resulting in alterations in synaptic function and cell-cell interaction pathways. A) Schematic representation of the experimental design utilized in this study and social interaction test delineates mice exposed to chronic social defeat into two groups depending on their SI: RS with SI above one, and SS with SI below one. CTR mice not exposed to stress exhibit an array of SI. Kruskal-Wallis One-Way ANOVA with multiple comparisons nonparametric test was used; CTR vs. RS p=0.0084, RS vs. SS and CTR vs. SS p<0.0001. B) PCA of the samples from the PFC shows clustering of DEGs, where CTR and RS cluster together distinctly from SS cluster. C) Heatmap of Top 100 DEGs showing majority are upregulated in SS mice, with preponderance in genes encoding proteins involved in the formation and regulation of the extracellular matrix scaffold. D) Normalized enrichment score (NES) of selected gene ontology (GO) pathways. E) ECMs’ number of fragments per kilobase of transcript per million mapped reads (fpkm) in astrocyte-specific IP vs. UB fraction for selected ECM gene transcripts point to changes in SS mice specific to astrocytes. *Sparc*: CTR IP vs CTR UB p=0.0383, CTR IP vs SS IP p<0.0001, SS IP vs SS UB p<0.0001; *Sparcl1*: CTR IP vs CTR UB p<0.0001, CTR IP vs SS IP p<0.0001, SS IP vs SS UB p<0.0001; *Ncan*: CTR IP vs SS IP p<0.0001, SS IP vs SS UB p<0.0001; *Vtn*: CTR IP vs SS IP p=0.0027, SS IP vs SS UB p=0.0318; *Lamb2*: CTR IP vs CTR UB p=0.013, CTR IP vs SS IP p<0.0001, SS IP vs SS UB p<0.0001, CTR UB vs. SS UB p< 0.0001; *Col1a2*: CTR IP vs SS IP p=0.0007, SS IP vs SS UB p=0.0005; *Col11a2*: CTR IP vs SS IP p=0.001, SS IP vs SS UB p=0.0002, *Col23a1*: CTR IP vs SS IP p=0.0001, SS IP vs SS UB p=0.0002, CTR UB vs. SS UB p = ns; *Col5a1*: CTR IP vs SS IP p=0.0169; *Thbs2*: CTR IP vs CTR UB p=0.001, CTR IP vs SS IP p=0.0183.

Previous studies reported that mice subjected to chronic social defeat exhibit short- and long-term behavioral changes ^46^. Principal Component Analysis (PCA) of mRNA expression data showed that gene expression in hippocampal and striatal astrocytes does not vary significantly among CTR, RS, and SS animals at 2-, 7- or 28-days post-stress (Fig. 1B, S1A, B). However, PFC astrocytes from SS mice have a gene expression profile that differs from CTR and RS astrocytes already at 2 days post-stress (Fig. 1B). We decided, therefore, to focus our studies on the PFC. Differential gene expression (DGE) analysis revealed 4205 and 3800 differentially expressed genes (DEGs) in astrocytes between SS and CTR or RS, respectively (p<0.05 and false discovery rate/adjusted FDR/padj<0.5). Only one gene was differentially expressed between RS and CTR mice (Table S1).

Hierarchical clustering allowed us to compile a list of gene expression changes between SS and CTR or RS in PFC astrocytes ranked by statistical significance (Fig. 1C, full list in Table S2). Of the top 100 ranked genes, 91 are upregulated in SS vs. CTR or RS PFC astrocytes (astrocyte enriched Table S3). Among these genes, some have previously been found to be enriched in reactive astrocytes, such as *Lgals1*, *Cd44*, *S1pr3*, *Lcn2*, and *Serpina3n* ^47^, suggesting that this subtype of astrocytes is induced under stress conditions in susceptible mice. We also found that the glutamate transporters *Slc1a2* and *Slc1a3*, glutamate synthetase (*Glul*), an enzyme involved in the conversion of glutamate to glutamine, and the gene *Kcnj10* that encodes the potassium transporter protein Kir4.1, are differentially expressed in SS mice, consistent with previous studies demonstrating a role for glutamatergic signaling in depressive-like behavior and MDD ^13,14,23,48,49^. Strikingly, and consistent with studies of other MDD models, ^25,50^ 36 of the top 100 DEGs encode extracellular matrix (ECM) proteins. These include upregulated genes encoding *Sparc* and collagen isoforms (*Col1a2*, *Col11a1*, *Col5a1*, *Col6a*, *Col8a2*, *Col23a2*), *Lamb2*, *Ncan*, *Vcan*, and *Fn1* (Fig 1C). Western Blot (WB) analysis from PFC lysates revealed differences in protein levels of SS and CTR mice (Fig. S1C, D), supporting functional significance. Gene ontology (GO) analysis also revealed over-representation of pathways related to collagen and ECM, as well as pathways related to synapse function, such as presynaptic and postsynaptic membrane complexes, ion channel complexes, or glutamatergic synapse among DEGs, raising the possibility that physical changes to synapses may take place during social-defeat-induced stress (Fig. 1D, Table S1). Considering that many of the astrocytes’ DEGs may also be expressed in other brain cell types, we asked whether these differences in gene expression were shared among non-astrocytic cells. To this end, we compared the profiles from SS and CTR of the PFC astrocytes mRNA (immunoprecipitated fraction-IP, astrocyte-specific) with the profile from all other PFC cell types (unbound fraction-UB, bulk mRNA, without astrocytes’ mRNA). Our results showed that the DEGs in PFC astrocytes are specifically enriched in SS IP fraction compared to UB fraction (Fig 1E). Thus, although DEGs are expressed in many cell types, these gene expression changes occur primarily in astrocytes.

### Synapse function and morphology are altered in stress susceptible mice

Because ECM proteins compose and regulate the extracellular space scaffold (ECS) ^43,44,51,52^ and are involved in synapse development and function, ^38–40,53^, we evaluated whether synaptic function is affected in SS mice. We performed whole-cell voltage-clamp recordings of AMPA-mediated excitatory postsynaptic currents (AMPA mPSCs) in SS and CTR animals. As shown in Fig. 2A, AMPA mPSCs amplitude, but not frequency (Fig. S2A), is significantly decreased in layer V pyramidal neurons of the PFC in SS mice. Furthermore, WB analysis of synaptosomal fractions isolated from the PFC of SS and CTR mice revealed no changes in AMPA glutamatergic receptors A1 and A2 (GluA1 and Glu A2) levels in SS mice (Fig 2B, Fig S2B). Together, these results suggest that while AMPA receptor localization to postsynaptic membranes is likely unaffected by stress exposure, synaptic function is perturbed. This led us to hypothesize that synaptic morphology might be a source of the changes in the AMPA postsynaptic currents.

**Figure 2.**
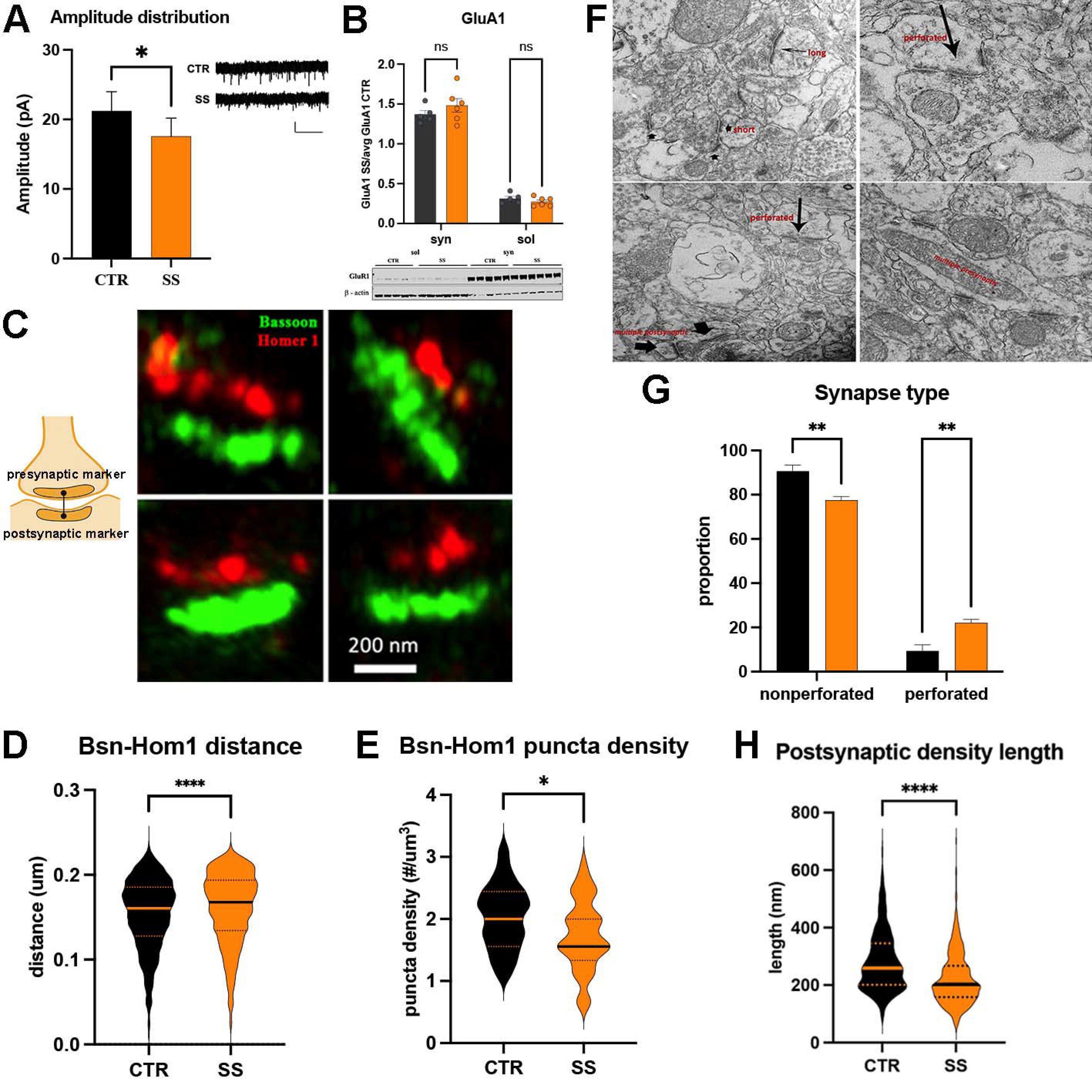
Synapse remodeling in stress susceptible mice. A) Amplitude of the excitatory postsynaptic currents is decreased in SS mice. B) WB analysis and representative blot showed no changes in the GluA1 protein enrichment in the synaptosomal fraction. C) Schematic diagram of method to measure the size of the synaptic cleft by imaging the distance between presynaptic and postsynaptic marker. Representative image of immunolabeling and expansion microscopy of the presynaptic (Bassoon/Bsn) and postsynaptic (Homer1/Hom1) markers. D) Bsn-Hom1 distance is significantly increased (*p*< 0.0001) in SS mice compared to CTR. E) The number of Bsn-Hom1 puncta are significantly decreased in the PFC of the SS mice (*p=0.0107*). F) Representative images from the prelimbic area of the PFC show synaptic structures that were quantified in G-H. G) Proportion of non-perforated synapses is significantly decreased and perforated synapses is significantly increased in SS mice (nonperforated in CTR vs nonperforated in SS p=0.0073; perforated in CTR vs perforated in SS p=0.0089). H) Length of the postsynaptic density is significantly decreased in SS mice (p<0.0001).

To assess changes in synapse morphology, we evaluated synaptic cleft size using expansion microscopy (Fig. 2C, D) ^54,55^. Quantitative measurement of distances between presynaptic (Bassoon, green) and postsynaptic (Homer1, red) scaffold proteins showed significantly greater separation in SS mice (Fig. 2D), from d=153+/- 42.1nm in CTR (n = 4; syn # = 5788) to d = 160+/- 42.9 nm in SS mice (n = 4; syn# = 5485), suggesting wider synaptic clefts. We also noticed that the number of bassoon-homer1 puncta is significantly lower in SS mice (Fig. 2E). Electron micrographs revealed that postsynaptic density length is significantly decreased in SS mice (Fig. 2H) and that more postsynaptic densities in these animals have a perforated appearance (Fig. 2F-G), a feature of synaptic plasticity also detected in other stress-induced settings ^56,57^. Taken together, and consistent with other reports, ^58–60^ our data suggest that chronic social stress induces morphological and functional synaptic changes.

### SPARC interacts with synaptic cleft proteins and its expression is induced in depressed mice

Our data showed that *Sparc* mRNA levels are the most significantly enriched among all the ECM encoding genes in SS mice (Fig. 1C), and we confirmed mRNA enrichement in SS PFC by RNAscope (Fig. 3A1-3), and SPARC protein enrichment in whole PFC tissue lysate (Fig. S1C). In SS PFC this enrichment is accompanied by decreased levels of *Klf4*, a transcription factor that inhibits *Sparc* expression, ^61^ (Table S1). In the adult brain, *Sparc* is abundantly expressed in astrocytes,^62^ and is translated in astrocyte processes surrounding synapses ^63^, accompanied by decreased levels of *Klf4*, a transcription factor that inhibits *Sparc* expression, ^61^ (Table S1). SPARC protein inhibits synapse formation and maturation by interacting with integrin b3 (ITGB3), preventing AMPA receptor accumulation at postsynaptic membranes ^38,39,64^. Outside of the CNS, SPARC promotes extracellular scaffold assembly and remodeling, and together with fibronectin and laminin, and binds collagens, proteoglycans, and hyaluronic acid together ^51^. Taking into account reported Sparc in the brain and periphery we hypothesized that in SS mice Sparc can affect synapse function in two ways (Fig. 3B): **1)** it can change the synapse function by modulating the postsynaptic membrane AMPA receptors, via interaction with the integrin b3 (ITGB3) and changes in membrane localization of these receptors affecting the glutamatergic signaling, similarly to what was reported in brain development ^38,39,64^, and **2)** in agreement with the reports of SPARC function in the extracellular scaffold assembly, we hypothesized that Sparc and ECMs in the top 100 DEGs in SS mice, interact to increase synaptic cleft space thus modulating glutamate accessibility in the synapse (Fig. 3B).

**Figure 3.**
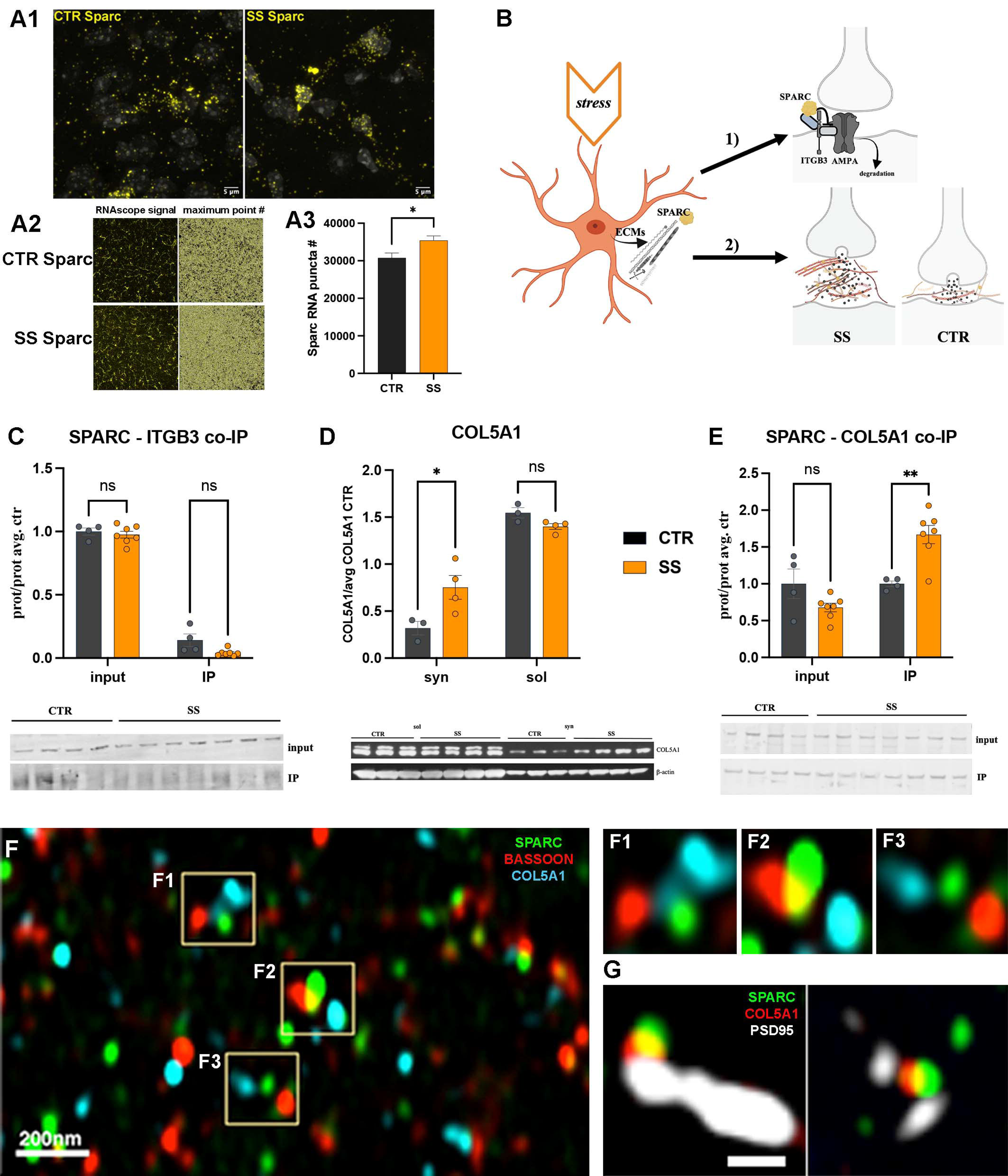
Enrichment and increased interaction between SPARC and COL5A1 corelates with increased synaptic cleft size. A) Representative low magnification image of the RNAscope analysis using *Sparc* probe (yellow, A1) of CTR and SS mice. A2) Representative images of puncta analysis, and number of puncta quantification (A3), shows increase in the PFC of SS mice (p=0.0345). B) Two possible hypotheses of the astrocyte function in stress induced depression were tested: 1) changes in synaptic function are a result of SPARC interactions with ITGB3, that disrupts ITGB3-AMPAR interaction and reduces the localization of AMPAR on the postsynaptic membrane; and 2) increased abundance of SPARC and ECMs increases their interactions and synaptic cleft size. C) WB quantification and blot image (below) show no changes in co-immunoprecipitation of ITGB3 with the SPARC antibody as bait in SS mice compared to the CTR mice (p=0.9254 for the input and p=0.09 for the IP). D) WB analysis of the distribution of COL5A1 protein in the synaptosomal and soluble fractions shows significant enrichment in the synaptosome in SS mice compared to CTR synaptosome fraction (p=0.0227), but no difference between CTR and SS soluble fractions (p=0.6444). E) co-IP analysis of the interaction between two ECMs SPARC and COL5A1 show increased interaction is SS mice (p=0.0043). F-G) Immunolabeling and expansion microscopy show colocalization of SPARC (green puncta) and COL5A1 (blue puncta) with the presynaptic marker Bassoon (F, red), and postsynaptic marker PSD95 (G, white).

We therefore aimed to investigate these hypotheses, by firstly checking if the interactions of SPARC with ITGB3 are increased in SS PFC by co-immunoprecipitation (co-IP). We confirmed that SPARC binds to ITGB3, but this interaction was unchanged in SS mice (Fig. 3C), and ITGB3 enrichment was also unchanged in SS mice (Fig. S3A). In addition, we evaluated SPARC interaction with ITGB1, another component of the ITGB3-AMPA receptor complex, and found this interaction to be unchanged in SS mice (Fig S3B). Thus, SPARC-integrin interactions are unlikely to explain synaptic changes in response to chronic-defeat-induced stress, suggesting that SPARC interaction with ECMs may modulate the synaptic function. To find the which ECMs from the top 100DEGs list are enriched in the synaptic fraction, we performed the WB analysis of CTR and SS PFC synaptosomes. We utilized an *in-silico* analysis HumanBase, a machine learning algorithm that uses genomic data to predict interactions between genes ^65,66^, to generate the SPARC interactome. This analysis showed multiple ECMs from the top100 DEGs in SS mice, including FN1, LAMC1, LAMB2, VCAN, and multiple collagen genes, COL5A1, COL4A1, COL4A2, to interact with SPARC (Fig. S3C). To evaluate the protein enrichment in synaptosomes we fractionated the whole tissue lysates and did WB with synaptosomal (syn) and soluble (sol) fractions with the antibodies against ECMs that interact with SPARC *in silico* and are enriched in our top DEGs. We found several collagens were present in the synaptosomal fraction, including COL5A1 (Fig. 3D), COL1A2, COL11A2, and COL23A2 (Fig. S3D1-3), but only COL5A1, although being enriched in the soluble fraction, is specifically enriched in the synaptosomal but not soluble fraction in the SS mice (Fig. 3D). Next, to determine if SPARC interacts with COL5A1, and if this interaction is changed in SS mice, we performed co-IP experiments. Remarkably, co-IP showed increased SPARC-COL5A1 interaction in these mice (Fig. 3E), but no increase in the interaction between SPARC and another synaptosomal enriched collagen, COL1A2 (Fig. S3E1). Because collagens are known to interact with integrins we aimed to evaluate if COL5A1 may interact with the ITGB3 and thus affect the AMPA-ITGB3 complex. We performed co-IP with COL5A1 as a bait and found no increase in the interactions between COL5A1 and ITGB3 or ITGB1 (Fig. S3F1-2). Next, by immunolabeling of PFC tissue we confirmed that SPARC and COL5A1 are co-localized with the synaptic markers, bassoon (Fig. 3F1-3) and PSD95 (Fig. 3G). Taken together, these experiments suggest that synaptic ECM changes involving increased binding of SPARC to COL5A1, localized in the synapse, may promote the alterations in synapse structure in SS mice.

### Sparc upregulation averts depressive-like behaviors in Sparc-KO mice

Because our data showed elevated gene expression and protein levels of SPARC in the SS mice, while RS mice show similar SPARC levels as controls, we aimed to examine if knockdown of SPARC could reverse the depressive-like phenotype. Evidence of the SPARC’s effect on behavior comes from the tests done on full body knockout mice. They are anxious but exhibit significantly less despair than the wild type (WT) mice ^67^. Thus, we sought to examine the effect of region-specific downregulation of *Sparc* on depressive-like phenotype. This was achieved in WT mice by injecting the prelimbic region (PrL) of the PFC with viral vectors carrying either scrambled control (scrm) small hairpin RNA (shRNA) or *Sparc* shRNA. The effect of *Sparc* downregulation was assessed by the WB analysis on the whole tissue homogenates as well as synaptosomal and soluble fractions (Fig. 4A1-2). We found a significant reduction of the SPARC protein in the PFC of *Sparc-*silenced mice compared to controls, and this did not change when mice were exposed to stress. SPARC was still detected in synaptosomes compared to soluble fraction, but this enrichment was significantly reduced in *Sparc-*silenced mice (Fig. 4A2). The effect on the behavioral phenotype was evaluated by utilizing a battery of behavioral tests, to assess social interaction, anxiety behavior in the open field (OF), and anhedonia-like behavior on the novelty suppressed feeding (NSF) before and after the chronic social defeat stress paradigm (Fig. 4B1-3). When mice were exposed to social defeat, both control and *Sparc-*silenced mice showed significantly lower social interaction compared to scrm injected controls (Fig. 4B1). Strikingly, in shRNA injected mice, exposure to social defeat resulted in deaths, that was absent in scrm treated mice. In addition, *Sparc-*silenced mice exposed to social defeat showed consistently higher latency to feed compared to scrm controls (Fig. 4B2). Control and *Sparc-*silenced mice showed similar behavior in the open field test, but *Sparc*-silenced mice exposed to stress exhibited significantly reduced time spend in the center of the open field compared to scrm controls (Fig. 4B3). Given that the NSF and OF tests is thought to tap into anxiety-like behavior, this result was in line with Sparc-KO behaviors reported by others. Importantly, these experiments showed that downregulation of *Sparc* expression does not rescue the MDD-like behavioral phenotype.

**Figure 4.**
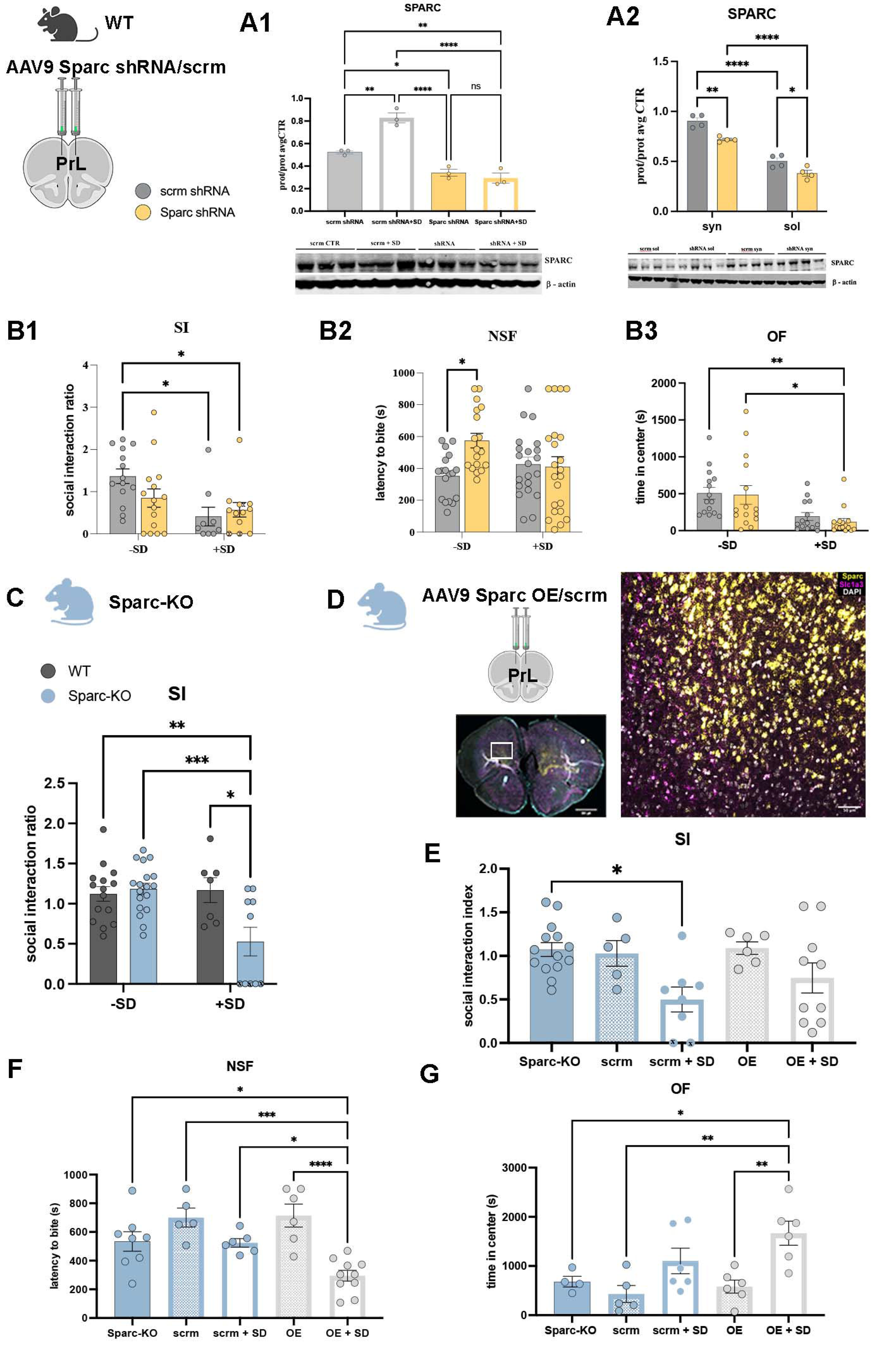
Increased expression of *Sparc* rescues the depressive-like phenotype in mice exposed to stress. A) Schematic diagram of region-specific downregulation of *Sparc* expression via injections of the AAV9 viral particles carrying scrm (control) and *Sparc* shRNA constructs. A1-2) WB analysis of the viral downregulation of *Sparc* expression in WT mice showed that SPARC protein enrichment in the PFC of WT mice injected with shRNA is not changed after exposure to social defeat (A1). The SPARC protein is enriched in the synaptosomal fraction of the PFC (A2), in scrm control and shRNA mice, but the shRNA mice have significantly reduced enrichment in both fractions. B1-3) Behavioral outcome of *Sparc* downregulation showed that Sparc shRNA injected mice have significantly decreased social interaction (B1, scrm -SD vs scrm +SD p=0.0138; scrm-SD vs *Sparc* shRNA+SD p=0.0303), increased latency to bite (B2, scrm -SD vs shRNA -SD p=0.0189), as well as decreased time spent in the center of the arena (B3, scrm -SD vs. shRNA +SD p=0.095, shRNA - SD vs. shRNA +SD, p=0.0159) for shRNA injected mice. C) *Sparc*-KO mice exhibit significantly reduced social interaction after exposure to social defeat (WT vs Sparc-KO+SD, p=0.0034; Sparc-KO -SD vs. Sparc-KO +SD, p=0.0007; WT+SD vs. Sparc-KO + SD, p=0.0110). D) RNAscope with the mouse-specific *Sparc* RNA probe shows that Sparc expression (yellow) level is restored in the PrL cortex of Sparc-KO mice. E) Social interaction after defeat stress is modestly improved by Sparc OE virus injection into the PFC of the *Sparc*-KO mice, but it does not reach statistical significance. F) *Sparc*-KO mice with restored *Sparc* expression showed statistically significant reduction in latency to bite after social defeat compared to all other groups of mice (OE +SD vs. Sparc-KO p=0.0186, vs. scrm p=0.0003, vs. scrm +SD p=0.0477, vs. OE -SD p<0.0001). G) Mice with restored Sparc expression exposed to social defeat showed statistically significant increase in time spent in the center of the arena (OE + SD vs. Sparc-KO p=0.0320, vs. scrm p=0.0027, vs. OE p=0.006).

To investigate the physiological effect of *Sparc* downregulation and subsequent exposure to social defeat stress on AMPA-mediated synaptic transmission we performed whole cell patch-clamp recordings of layer V pyramidal neurons in acute PFC slices (Fig. S4A1-4). In mice injected with the scrm control virus, social defeat stress led to a significant decrease in the amplitude but not the frequency of AMPA mPSCs. In mice where *Sparc* was silenced, the baseline amplitude of AMPA mPSCs was decreased compared to control mice, but stress did not decrease it further. On the other hand, the frequency of AMPA mPSCs, not changed by the mere silencing of *Sparc*, was significantly decreased upon social defeat stress in *Sparc-* silenced mice compared to control. These results suggest that *Sparc* downregulation with or without stress does not affect postsynaptic component of glutamatergic synapses and resulting mPSCs. Furthermore, WB analysis of AMPA and NMDA receptors subunits localization showed that these were not significantly changed after *Sparc* downregulation (Fig. S4B1-4). This further confirmed that the effects of SPARC on stress-induced synaptic transmission changes are not mediated by glutamatergic receptor localization/enrichment on the postsynaptic membrane, unlike what has been shown previously for developing synapses ^38^. Data from these experiments further indicate that downregulation of *Sparc* does not reverse depressive-like phenotype after exposure to chronic social defeat, and that this is not a result of the changes in the localization of the AMPA glutamatergic receptors in synaptosomes. The amplitude distribution and frequency are still affected by *Sparc* perturbations, pointing to a different mechanism of action, one in which SPARC may modulate the extracellular space architecture in the synapses.

Since *Sparc* downregulation failed to rescue the depressive phenotype, we sought to examine the effect of its upregulation in the PFC. For this purpose, we used CRISPR to generate SPARC full body knockout mice (Sparc-KO) and observed that they have significantly decreased social interaction after social defeat (Fig. 4C). Similarly to the mice with Sparc downregulation, Sparc-KO mice exposed to stress also died in the course of social defeat. To confirm that Sparc is beneficial for positive response to social defeat stress we sought to restore Sparc expression in the PFC. This was achieved by injection of the overexpressing (OE) Sparc AAV9 viral vector or scrm control into the PrL area of Sparc-KO mice (Fig. 4D). Behavioral effects were accessed with SI, OF, and NSF tests. Unlike control mice, *Sparc* overexpressing mice did not show decreased social interaction after social defeat, while scrm mice did (Fig. 4E). Importantly, mice injected with OE also had a significant decrease in the latency to bite after social defeat (Fig. 4F) and increased time spent in the center of the arena (Fig 4G), clearly showing a positive behavioral outcome after *Sparc* expression was restored in the PrL of Sparc-KO mice.

Taken together, these data suggested that in chronic social stress astrocytes increase expression of the ECM proteins, including SPARC and collagens, followed up by their greater interaction. This in turn coincides with the remodeling of the synapse morphology, including synaptic cleft height and protein content, prompting changes in the synaptic activity. Notably these changes appear to be beneficial, since a full body *Sparc* knock out mice, but with virally induced expression in the PFC, exhibit significantly reduced anxiety and increased social interaction after exposure to stress.

## Discussion

Alterations in synaptic morphology, density, and excitatory transmission are hallmarks of major depressive disorder in humans and animal models ^4,6,16,68–72^. MDD patients show fewer synapses, and decreased expression of synapse-related genes in the prelimbic cortex^58^, similarly to chronic stress-induced reduction in the number of synapses and spines^59,73^. By implementing a chronic social defeat stress model of MDD, we uncover that these synaptic changes are astrocyte dependent and associated with alterations in synapse morphology.

To the best of our knowledge, the role of ECM, especially Sparc and collagens, in the modulation of the synapse morphology and activity in stress induced MDD and its mechanism of action have not been reported previously.

Glutamatergic transmission depends on the molecular and morphological architecture of the synapse, including the size of the synaptic cleft, and the speed of glutamate diffusion through the cleft, which is dependent on the interstitial space viscosity and charge of the proteins ^41,74,75^. It has been shown that extracellular matrix fibrils form a net within the synaptic cleft ^76^ that can determine the distance between the pre and post synaptic buttons. A narrower cleft leads to a higher concentration of neurotransmitters and enhanced activation of receptors ^42^. In addition, the molecular composition of the cleft has a significant effect on the diffusion of glutamate, where such ECMs, like negatively charged glycosaminoglycans, may affect the diffusion of negatively charged glutamate^77^. The ECMs may also increase the interstitial viscosity and may regulate the size of the extracellular space ^41^. Here, we have shown that upregulation of genes encoding ECMs in astrocytes is associated with their protein enrichment in the synaptosomal or soluble cell fractions. This is accompanied by increased protein-protein interaction, specially of SPARC with COL5A1, in stress susceptible mice. *Sparc*, the gene with the highest upregulation in SS mice, participates in the extracellular scaffold remodeling, serving as a matricellular chaperone mediating the ECM disassembly and degradation of the scaffold networks ^78,79^, as well as inhibiting degradation and disassembly of already present collagen fibrils ^80^. Interestingly it has been shown that SPARC serves as a chaperone molecule that organizes collagen fibrils into larger diameter fibers in the dermis ^78,81^. This suggests that increased SPARC-COL5A1 interaction increases the collagen fibers content in proximity and within the synaptic cleft, where we found the significantly higher enrichment of COL5A1, and which may extend the height of the synaptic cleft observed in SS mice. Interestingly, mutations in *Col5a1* are causing subtypes of Ehlers-Danlos syndrome, a connective tissue disorder in which patients also experience psychiatric symptoms, including depression ^82,83^.

In addition, it reasonable to speculate that the additional ECMs secreted into extracellular space would increase its viscosity, reducing the diffusion of glutamate and ions. This may impede postsynaptic activity even without changes in AMPA and NMDA receptor composition on the postsynaptic membrane, similar to what was proposed previously ^84^. This would explain the changes in AMPA mEPSCs in the absence of increased interaction between the SPARC and ITGB3-AMPA complex, described in the developing brain ^38,64^. Indeed, our data showing that the amplitude, but not the frequency of postsynaptic electrical activity is affected, further supports the role of the increased size and viscosity of the synaptic cleft.

Moreover, it is possible that SPARC influences these changes via ECM scaffold remodeling in perisynaptic areas, and in the extracellular space between astrocyte and neuronal membranes, supported by the finding of an increased interaction of SPARC and COL23A1 outside of the synaptosome. These data and the literature would point to the involvement of ECM in reshuffling of the protein interactions at the membranes, including in the synaptosome and in the extracellular space between the cells. It is conceivable that the altered extracellular scaffold changes cell-cell interaction between astrocytes and neurons, or astrocytes themselves, disrupting the equilibrium of cell adhesion and cell-cell communication.

We have also found a reduction in the number of synapses and the length of postsynaptic density in SS mice. It is noteworthy that the number of perforated synapses almost tripled in the deep layers of the prelimbic cortex in stress susceptible mice. This is similar to previously reported effects in the infralimbic area of the PFC of the rats exposed to chronic stress ^59^. Perforated synapses contain multiple postsynaptic densities and high densities of the AMPA receptors, suggesting increased synaptic efficacy (reviewed in ^85,86^). Interestingly, it has been shown that SPARC has a role in the elimination of cholinergic synapses, characterized by the presence of SPARC-dependent pre- and postsynaptic membrane invaginations ^87^. In the context of our findings, this would suggest that SPARC expression induced by stress may be responsible for changes in synapse morphology, including increased numbers of synaptic invaginations, and perforated synapses. Thus, the effect of SPARC on synapse function is two tiered: boosting the buildup of the ECM scaffold and modulation of the synaptic membrane morphology.

The behavioral effect of ECM dependent synapse remodeling is less clear. When *Sparc* was silenced specifically in the PrL, and thus effectively blocked from its upregulation in mice exposed to chronic social defeat, these mice exhibited a depressive-like phenotype, same as the wild type mice with upregulated *Sparc* after social defeat. Strikingly, overexpression of SPARC in the PFC of *Sparc*-KO mice reversed this phenotype, pointing to the important role of this protein in the stress response related to PFC. To understand whether upregulation of ECM genes have a positive or detrimental long-term effect on the neuronal function in stress warrants further examination. For example, in this study, restored expression of *Sparc* appears to promote better behavioral outcomes in mice lacking *Sparc*, at least for a short period after exposure to social defeat stress.

In summary, we have shown the role of astrocytes in regulating synaptic morphology, restructuring of the synapse, and reducing the firing amplitude of the pyramidal neurons in the prefrontal cortex. These events are a result of the astrocyte specific increase in the ECMs expression, increased interaction of SPARC and fibrillary collagens, and the resulting increase of synaptic cleft heigh. These changes are associated with the changes in synaptic activity linked to the increased size and possibly viscosity of the synaptic cleft. This study points to a complex nature of astrocyte-neuronal interaction and highlights the importance of examining multiple cell types and their role in the neuropathological processes, as well as circuitries regulating behavior. Our data points to the significance of the astrocytes’ secreted proteins and their role in the modulation of the synaptic cleft architecture that determines optimal synaptic activity. The identification of additional cell specific signatures will further our understanding of the onset and progression of the neurological diseases, including MDD, and facilitate the search for novel therapies and protocols for precision medicine.

## Methods

### Mice

We utilized astrocyte specific TRAP mouse line in which L10a-GFP is expressed under the astrocyte specific Aldh1l1 promotor ^45^. The mice were group housed in conventional rooms in the Rockefeller University animal facility in a 12-hour light/dark cycle and access to food and water ad libitum. For tissue harvesting mice were anesthetized with CO2, decapitated, and brain tissue was dissected. For immunocytochemical, protein, and electron microscopy analyses we used C57Bl/6J mice purchased at 6 weeks of age from the Jackson Laboratory. Mice were left to acclimatize to the homeroom for at least one week prior to the commencement of experiments. All procedures and animal care were approved by the Rockefeller University IACUC.

### *Sparc* knockout mice

This knockout was generated by the Rockefeller University CRISPR and Genome Editing Resource Center at the Transgenic Mouse Core. Target sequence of guide RNAs used for editing was Sparc-crRNA-B 86 : GTGTGGGTCCTCAGATCGTC *AGG* and Sparc-crRNA-D-72 ; ACAGAGGTCATGGTCACATC *CGG*.

Editing template was lssDNA amplified from plasmid using primers Sparc-ss-F1(GCAAGATGGTTTGGTGGAAGA) and Sparc-ss-R2(GCTGTACAACCATGAAGACGG). Mice were genotyped with primer set Sparc-Scr-F1 (GCTTCCAGGCACCTAGTACA)+Sparc-Scr-R2(CCTTCTCGACTGAATCCCCA) until homozygous mice were obtained (∼F6 generation). Absence of SPARC protein in the brain was also confirmed by WB with anti-Sparc antibodies (LSBio, R&D, Novus), and absence of mRNA by the RNAScope with specific mouse mRNA probe.

### TRAP mRNA isolation

Immunoprecipitation and RNA isolation was performed as previously described ^45,88^. Briefly, the relevant area is dissected and minimum of three replicate samples per brain region isolated; for nucleus accumbens individual mice were pooled into one sample. Immunoprecipitation was performed overnight with the mixture of two monoclonal anti-GFP antibodies. RNA was purified using Quiagen Rneasy Micro Plus kit, following the manufacturer’s protocol, and quantity and quality determined with a Nanodrop Spectrophotometer and Agilent Bioanalyser. For each sample, RNA was amplified using the Ovation RNA-seq System V2 (NuGEN). Library preparation and amplification was done by the Rockefeller University Genomic Facility, and Illumina HiSeq platform was used.

### Bioinformatics analysis

We have used the fastp ^89^ with default parameters for the quality control of raw reads. Raw reads were mapped to the mouse mm10 genome by using STAR ^90^ in default parameters. The systemPipeR ^91^ were invoked to build the workflow of calculating the RPKM and detecting the Different Expressed genes. R package DESeq2 were used for the detection of the differential expressed genes ^92,93^ and the cutoff for the DEGs is FDR < 0.05 and log2 folder change > 2.

### Surgeries and viral injections

Wild type C57Bl/6J and Adlh1l1-TRAP mice underwent stereotaxic surgeries under general ketamine-xylazine anesthesia. Adeno-associated viral particles, serotype AAV9 carrying the shRNA vector against mouse *Sparc* (cat # shAAV-272934) or scrambled control (cat # 1122) shRNA were purchased from Vector Biolabs. For surgeries, mice were placed in the stereotaxic apparatus and the injection was guided by the AngleTwo program. Both sides of PFC were injected, and the injection coordinates were: +/-0.40mm lateral, 1.70mm anterior and -2.35mm ventral from the bregma. Each mouse was injected using the 10mL Hamilton syringe and 33g Hamilton needle aided by an infusion pump over 10 minutes (World Precision Instruments), with 10 minutes rest time after the infusion to allow the liquid to diffuse. Mice were kept group housed in a homeroom under close supervision for 3 weeks to allow recovery.

### Behavioral testing

#### Chronic social defeat

The social defeat stress was performed as previously described^28^. Briefly, Aldh1l1-TRAP and CDBL/6J mice were kept group housed minimum of two weeks prior the experiment. Mice were kept in a 12-hour light-dark cycle, at room temperature and with water and food provided *ad libitum*. Mice were randomly selected to control and experimental groups. Experimental group mice were housed with the aggressive mouse in a large cage with the Plexiglas divider, that allows visual and olfactory, but no physical contact. Direct contact with the CD1 mouse was allowed for 10 minutes daily, for ten days. Control mice were kept group housed, 2-3 mice per cage, in the same room. A day after the last day of social defeat, all mice underwent social interaction test.

#### Social interaction test (SI)

measured the time mouse spent close to a mouse in the arena that were separated by Plexiglas divide, as opposed to sulking around in corners. Mice were placed in stress susceptible group if they were prone to depression (i.e. avoided social interaction with novel mouse). Finally, we sacrificed and dissected prefrontal cortex brain area of both control and depression-susceptible mice and proceeded with IP and sequencing.

#### Novelty suppressed feeding (NSF)

Mice were food-deprived for 24 hours and then placed in an open field (50 x 50 x 22.5 cm) with a 1.5 cm long regular chow food pellet at the center, on the 10 cm diameter filter paper. The amount of time it takes for the mouse to take a bite of the food is recorded (latency to feed). The time to approach the center/filter paper, as well as time to sniff without biting, the number of sniffs, and the number of biting bouts were also recorded. Maximum time recording is 15 minutes.

#### Open field test (OF)

Mice were placed individually in plastic boxes and locomotor activity is recorded from above using video camera and Ethovision software. Movement is tracked for the duration of the test (1 hour) and can be broken down into shorter epochs, if necessary. After testing, mice were placed back into their home cage.

### Slice electrophysiology

For all patch-clamp experiments, we used 11–12-week-old wild-type mice, injected with the shRNA at 6 weeks, and subjected to chronic social defeat stress at 9-10 weeks. Coronal slices containing the mPFC (350 μm thickness) were cut at 2 °C using a Vibratome 1000 Plus (Leica Microsystems, USA) in a NMDG-containing cutting solution (in mM): 105 NMDG (N-Methyl-D-glucamine), 105 HCl, 2.5 KCl, 1.2 NaH2PO4, 26 NaHCO3, 25 Glucose, 10 MgSO4, 0.5 CaCl2, 5 L-Ascorbic Acid, 3 Sodium Pyruvate, 2 Thiourea (pH was around 7.4, with osmolarity of 295–305 mOsm). After cutting, we recovered the slices for 30–45 min at 35 °C and for 1 h at room temperature in the recording solution. The physiological extracellular solution used for recordings containing (mM): 125 NaCl, 25 NaHCO3, 25 glucose, 2.5 KCl, 1.25 NaH2PO4, 2 CaCl2, and 1 MgCl2 (bubbled with 95% O2 and 5% CO2). All experiments were performed on morphologically identified layer V pyramidal neurons with the help of a Multiclamp 700B amplifier (Axon Instruments, Molecular Devices, Sunnyvale CA, USA) and an upright BX51WI microscope (Olympus, Japan) equipped with Nomarski optics. AMPA-mediated excitatory postsynaptic currents (AMPA mPSCs) were studied by whole-cell voltage-clamp recordings in the presence of TTX (300 nM), AP-5 (10 μM), and Bicuculline (30 μM). For all experiments, we used an intracellular solution containing (in mM): 126 K gluconate, 4 NaCl, 1 MgSO4, 0.02 CaCl2, 0.1 BAPTA, 15 Glucose, 5 HEPES, 3 ATP, and 0.1 GTP. The pH was adjusted to 7.3 with KOH, and the osmolarity was adjusted to 290 mosmol l−o with sucrose. Patch electrodes, fabricated from thick borosilicate glasses, were pulled and fire-polished to a final resistance of 5–7 MΩ. Recordings with either leak currents >100 pA or a series resistance >15 MΩ were discarded. Action potential threshold was measured from the first action potential to avoid any confounding effects of adaptation. Data were acquired at a sampling frequency of 50 kHz and filtered at 1 kHz and analyzed offline using pClamp10 software (Molecular Devices, Sunnyvale, CA, USA). All electrophysiological data are expressed as means ± SEM.

### Expansion microscopy

Animals were deeply anesthetized with Nembutal and intracardially perfused with 4% PFA and brains dissected and cryopreserved in a series of sucrose solutions. After freezing, brains were sectioned at 40 micron-thick sections. Immunocytochemistry was performed as follows: after permeabilization and blocking for 1 hr in 4%BSA and 4% NGS in PBST, with 1/5 volume of DMSO to enhance the penetration of the antibodies, samples were incubated overnight at 4^0^C with the Bassoon and Homer1 antibodies (1/300 dilution) in 2%BSA and 2%NGS in PBST. Sections were washed and appropriate fluorescent secondary antibodies were applied. Brain slice expansion was performed as previously described ^54^. Briefly, tissues were treated with 100 times volume of a slice with the proteinase K (dilution 1/100) in digestion buffer (50mM Tris, 1mM EDTA, 0.5% TX-100, and 0.8M guanidine HCl) for 12 hrs. Next, slices were washed in double-distilled H2O for up to 2 hrs to let them expand. If the fluorescence appears bleached, the primary and secondary antibodies were applied again overnight at 4 °C. Upon completion of re-staining, gels were subsequently washed twice in PBS for 10 minutes to remove excess antibodies and fully expended in double-distilled H2O prior to imaging.

Expended tissues were imaged on an inverted Zeiss LSM 880 laser-scanning confocal microscope equipped with an Airyscan super-resolution module (Carl Zeiss AG, Germany). All images were acquired using a 63 × 1.4 NA oil objective lens (Plan-Apochromat, 1.40 Oil DIC M27), with the pinhole set at 1.25 AU. For imaging synaptic distance and density, Z-stack dimensions were set to 20 µm. Images were acquired as 868 × 868 pixels (zoom factor = 3.6) with a Z-interval of only 0.04 µm using piezo Z sectioning, corresponding to a voxel dimension of 0.04 × 0.04 × 0.04 µm in x, y, and z directions. To reduce overall photobleaching and crosstalk between fluorescent channels as much as possible, dual channel simultaneously scanning was adapted using MBS 488/561 as the dichroic beam splitter. Airyscan processing was done in 3D mode at default settings using Zeiss Zen 2 software (black edition).

Image analysis was performed using Imaris® (version 9.1.2) software (Bitplane Inc.) Pre- and post-synaptic puncta were detected as spots in Imaris, and the pre- and post-synaptic protein pairs was analyzed in Imaris XTension based on nearest neighbor pairs with cut-off at 250 nm for the identification of Bassoon - Homer1 synaptic pair.

### Transmission electron microscopy

Brains were perfused 2–3 min with saline and followed by fixation, containing 2% paraformaldehyde and 2.5% glutaraldehyde in 0.75 M sodium cacodylate buffer (pH 7.4) at 25 °C for 5 min. The brains were kept in the fixative solution at 4 °C overnight. Vibratome sections were generated, washed, post-fixed with 1% osmium tetra-oxide for 1 h, underwent en bloc staining with 2% uranyl acetate for30 min, dehydrated by a graded series of ethanol, removed ethanol with propylene oxide, infiltrated with a resin (EMBed 812; Electron Microscope Sciences, Hatfield, PA) and embedded with the resin. After polymerization at 60 °C for 48 h, ultra-thin sections underwent post-staining with 2% uranyl acetate and 1% lead citrate and were examined in the electron microscope (100CX JEOL, Tokyo, Japan) with the digital imaging system (XR41-C; Advantage Microscopy Technology (AMT) Corp., Denver, MA). Trypanosomes were cryoprotected with 20% bovine serum albumin (BSA) and applied to a high-pressure freezing procedure (EMPACT2, Leica Microsystem System, Wetzlar, Germany). Cells were transferred to a freeze substitution device (EM AFS2, Leica Microsystem System, Wetzlar, Germany), incubated with 0.2% Uranyl acetate in 95% acetone at -90 C°, and embed in Lowicryl HM20 at -35 C°. Ultrathin sections were cut, and grids were washed with water and uranyl acetate (1%), and lead citrate (1%) was applied. The sections were examined in the electron microscope (100CX JEOL, Tokyo, Japan) with the digital imaging system (XR41-C, AMT Imaging, Woburn, Massachusetts). The analysis was done on twenty non-overlapping EM fields from the deep layer of the prelimbic cortex, in four biological replicates from control and stress susceptible mice. The length of the postsynaptic density was measured using ImageJ software.

### Western Blot

After light CO2 anesthesia, mice were sacrificed by decapitation, and the pre-frontal cortex rapidly dissected. Tissue was sonicated for 20-30 secs in buffer containing complete protease and phosphatase inhibitors (7.5mL milliQ/ddH20, 1mL 10xPBS, 1.5mL 20% SDS).

The protein concentration of the tissue lysates was quantified by BCA Protein Assay (Bio-Rad). 12 µg-30 µg of proteins were mixed with the appropriate amount of 1X LDS Sample Buffer (Thermofisher), and heated for 5 min at 85°C. Then samples were loaded on a 4-12% Bis-Tris gel for electrophoresis and run at 200V for 55 minutes. Proteins were transferred at 35V for 2 hours onto a PVDF membrane, using XCell SureLock Mini-Cell Electrophoresis System (Thermofisher). The membrane was left to dry overnight at room temperature. The membrane was incubated with blocking buffer for 1 hour in agitation (Odyssey Blocking Buffer in TBS). Subsequently the membrane was incubated with indicated primary antibody, for 2 hours in agitation. Primary antibodies used were BLANK + anti-β-Actin (mouse, 1:7500, Cell Signaling) was used to normalize protein load. The membrane was washed 3 times with 1x TBS-T over the span of 15 minutes. The membrane was probed with either Goat anti-mouse HRP, Goat anti-Rabbit HRP, Rabbit anti-Goat HRP Secondary antibody for 1 hour in agitation (1:5000). The membrane was washed two times with 1X TBS-T for 10 minutes and once with 1X TBS for 8 minutes. Detection was carried out with PLUS Chemiluminescent Substrate (Thermofisher), based on the chemiluminescence of luminol. Quantitative densitometry of protein bands analysis was performed with ImageJ software. The protein enrichment was determined by calculating protein bend optical intensity over the average bend optical intensities of the control samples, all normalized by the respective loading control (actin/GAPDH) bends first.

### Synaptosome fractionation isolation

For synaptosome preparation, all steps were performed at 4°C or on ice, and all buffers contained protease and phosphatase inhibitors (Thermofisher). Mice were lightly anesthetized in a CO2 chamber and decapitated. Rapidly dissected PFC were homogenized in Syn-Per Reagent containing complete protease and phosphatase inhibitors (Thermofisher), ∼1 ml/100 mg tissue. Samples were centrifuged at 1200 × g for 10 min. The resulting supernatant was then centrifuged at 15,000 × g for 20 min. Pellets (synaptosomes) were resuspended in Syn-Per Reagent. Protein concentration was quantified by BCA Protein Assay (Bio-Rad) and samples were stored long-term at −80°C.

### Co-Immunoprecipitation

After light CO2 anesthesia, mice were sacrificed by decapitation, and the prefrontal cortex rapidly dissected. Tissue was mechanically homogenized in Co-IP Lysis Buffer (containing 1% NP-40), centrifuged at maximum speed for 10 minutes. Protein concentration of the resulting supernatant was quantified via BCA Protein Assay. The 50uL of Dynabeads per IP were resuspended via pipetting. The beads were washed 5X with PBS and supernatant was removed by placing the beads on the magnet/dynal. The beads were then resuspended in 200uL of AB binding and washing buffer and 5ug of SPARC primary antibody per IP sample (Novus Biologicals AF942). After incubating in this solution on a roller for 2 hours at room temperature, the supernatant was removed, and the beads were washed 5X with AB binding and washing buffer. The beads were then resuspended to their original volume in Co-IP Lysis Buffer. 50uL of beads + 200 uL of the protein extract (at 3ug/uL) were incubated in an orbital shaker overnight at 4°C. The supernatant was removed and kept as INPUT1. The beads were washed 5 times with 1X PBS, and the protein was eluted in 1X LDS Sample Loading Buffer for 10 min at 70 °C. These samples were placed on the dynal and loaded as INPUT2. INPUT1 and INPUT2 were both loaded onto a 4-12% Bis-Tris gel for electrophoresis and run at 200V for 55 minutes. Proteins were transferred at 35V for 2 hours onto a PVDF membrane, using XCell SureLock Mini-Cell Electrophoresis System (Thermofisher). The membrane was left to dry overnight at room temperature, followed by incubation with blocking buffer for 1 hour (Odyssey Blocking Buffer in TBS). Subsequently the membrane was incubated with indicated primary antibody, for 2 hours. The list of primary antibodies used is in Supplemental Table 3. Anti-β-Actin (mouse, 1:7500, Cell Signaling) was used to normalize protein load. The membrane was washed 3 times with 1x TBS-T over the span of 15 minutes. The membrane was probed with either Goat anti-mouse HRP, Goat anti-Rabbit HRP, Rabbit anti-Goat HRP Secondary antibody for 1 hour in agitation (1:5000). The membrane was washed two times with 1X TBS-T for 10 minutes and once with 1X TBS for 8 minutes. Detection was carried out with PLUS Chemiluminescent Substrate (Thermofisher), based on the chemiluminescence of luminol. Quantitative densitometry of protein bands analysis was performed with ImageJ software, and values were normalized for the actin input to enable the quantification of collagen antibody interacting with SPARC from the control and stressed mice.

### RNAscope

In situ RNA detection was performed with the Advanced Cell Diagnostic (Bio-Techne, MN) reagents, kits, and ACD-Boekel hybridization oven (cat # 321710). We used Multiplex Fluorescent Reagent Kit v2-Mm with TSA Vivid Dyes (cat # 323280) and specific probe for mouse Sparc (cat # 466781-C1). In situ was performed according to the manufacturer’s protocol. In short, 14-micron-thick cryostat sections from the mouse PFC were washed in PBS and fixed for 1 hr in ice-cold 4% PFA. Slides were then washed in a series of ethanol dilutions (50-100%) and treated with H2O2 for 10 mins. Target retrieval was done with the 1x Target solution heated to 96-98^0^C, for 10 mins, washed in H2O, 10% ethanol, and baked at 60^0^C for 5 mins. After overnight drying sections were treated with protease III at 40°C for 10 mins, and probe was applied for 2 hrs at 40°C. Slides were then washed in wash buffer, signal was amplified and Vivid dyes applied. Slides were counterstained with DAPI and coverslipped with ProLong Gold. Images were taken at the LSM900 Zeiss confocal microscope using same imaging parameters for all samples. Quantification of the RNA signal was done from these images using the maximum point number.

## Funding and acknowledgements

This work was supported by the NIDA P30 DA035756 Pilot Project Award and NIA 1RF1AG059770-01 (to A Milosevic).

We thank Dr. Nathaniel Heintz for a generous gift of the Aldh1l1-TRAP mice (project Gene Expression Nervous System Atlas (GENSAT) Project, NINDS Contracts N01NS02331 & HHSN271200723701C to The Rockefeller University). We are grateful to Dr. Cagla Eroglu (Duke U) and Dr. Corey Harwell (UCSF) who read the manuscript and offered thoughtful suggestions. We extend special thanks to Dr. James E. Goldman (Columbia U) for many valuable discussions on the role of astrocytes in the brain, and Dr Lisa Schimmenti (Mayo Clinic, MN) for illuminating discussion about Ehler Danlos syndrome and collagens. We would also like to acknowledge immense help from the Rockefeller University Resource centers, including Comparative Bioscience Center, Transgenic Center, Genomic center and Bioinformatics Center, as well as Bioimaging center at memorial Sloan Kettering Cancer center.

Role of Authors: all authors take responsibility for the accuracy of the data and analysis. All authors had full access to all the data in the study. Study concept and design: AM, SR. Acquisition of the data: AM, PRR, LM, JC, TF, PCDV, KM, FC, KC, LP. Analysis and interpretation of the data: AM, PRR, RZ, WW, LM, JPR, CM, SS, OT. Drafting of the paper: AM. Obtained funding: AM, RS. Technical support: WW, TF, PCDV, KM, CM, FC, KC, LP. Study supervisor: AM.

## Supplemental Figures

**Supplemental Fig 1.**
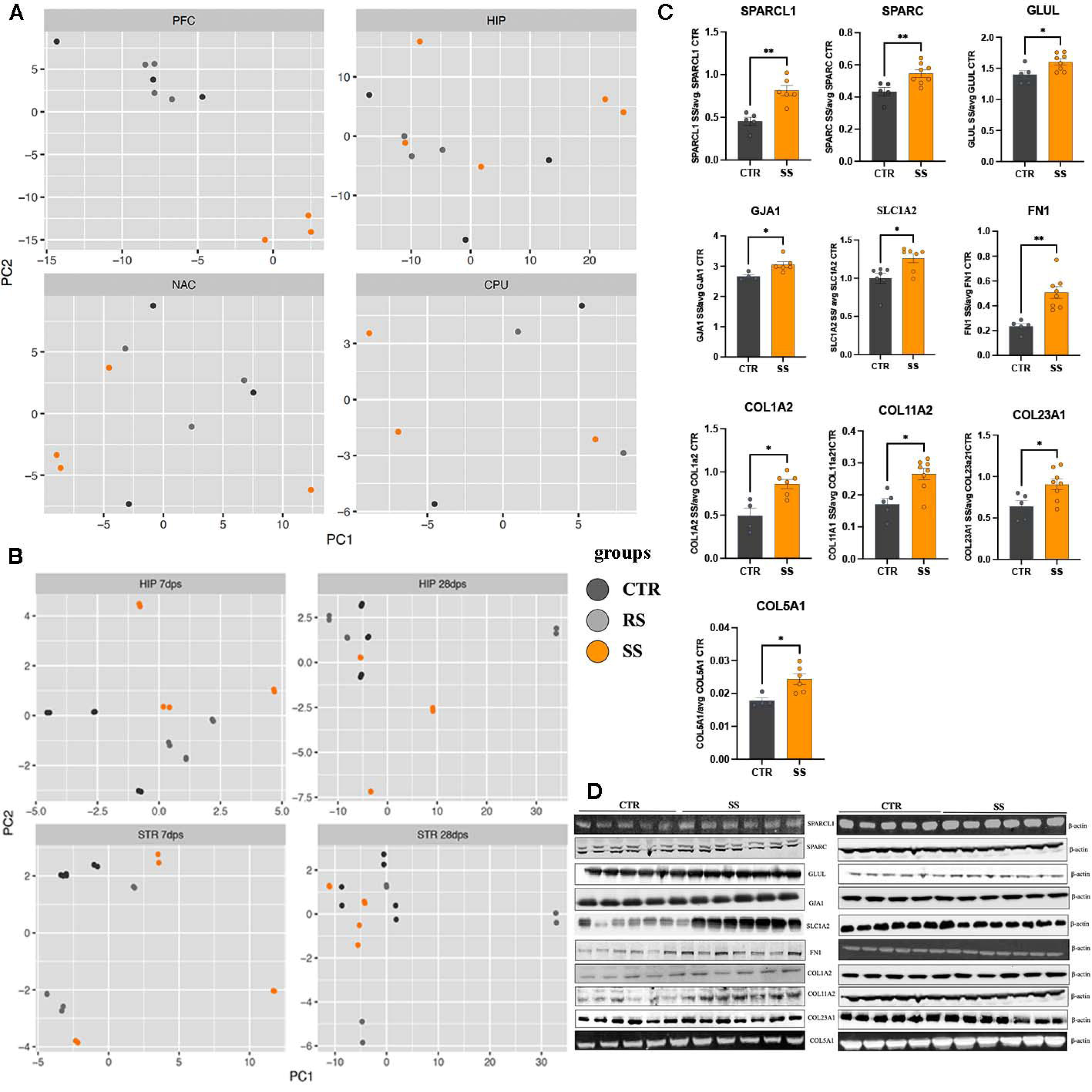
A) PCA analysis of the samples from the PFC, HIP, NAC, and CPU shows significant clustering of differentially expressed genes (DEGs), only in the PFC, but not other brain regions. B) PCA analysis of HIP and striatum (NAC and CPU) samples isolated from the brain of the CTR, RS and SS mice 7- and 28-days post stress did not show clustering of sample from the same experimental group. C-D) Protein enrichment graphs (C) and WBs blots (D) of the selected DEGs show statistically significant increase in the PFC of the SS mice. Differential enrichment in protein expression between CTR and SS were calculated with Mann Whitney unpaired t test, and p values were following: soluble protein acidic rich in cysteine like 1 (SPARCL1), p=0.0043), soluble protein acidic rich in cysteine (SPARC), p=0.0062), glutamate ammonia ligase (GLUL), p=0.0295; gap junction protein alpha 1 (GJA1), p=0.019; solute family carrier 1 member A2 (SLC1A2), p=0.0379, fibronectin 1 (FN1), p=0.0016; three fibrillar collagen type alpha chain proteins (COL1A2, COL11A2, COL5A1) are all enriched in SS (p=0.019, for COL1a2, and COL11A2, p=0.0381 for COL5A1). a nonfibrillar transmembrane collagen 23 alpha 1 chain (COL23A1), p=0.0295.

**Supplemental Figure 2.**
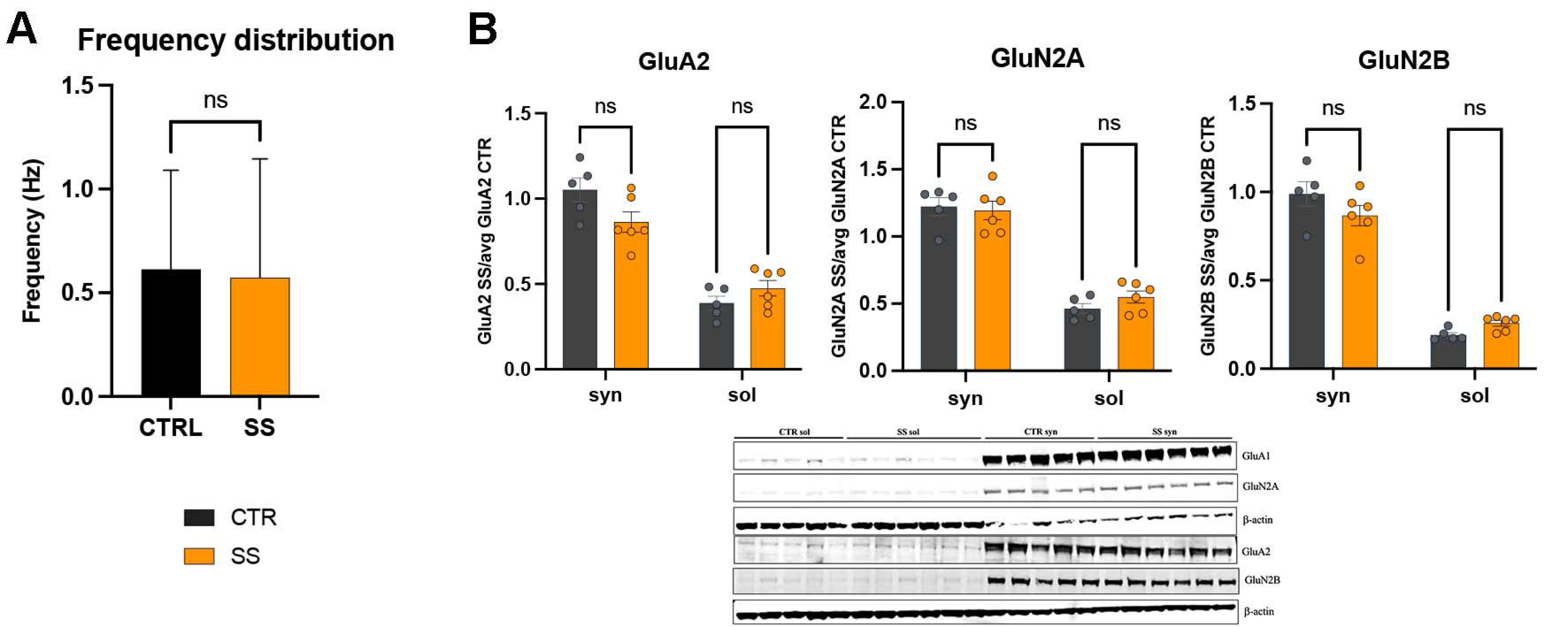
WB analysis of the protein enrichment in the synaptosomal and soluble fractions isolated from the PFC of SS and Ce mice. A) The frequency distribution of the mEPSCs is not changed between SS and CTRL mice. B) The protein enrichment of the AMPA glutamate receptor 2 (GluA2), and two NMDA receptors (GluN2A and GluN2B) are not significantly changed in either synaptosomal (syn) or soluble (sol) fractions.

**Supplemental Fig 3.**
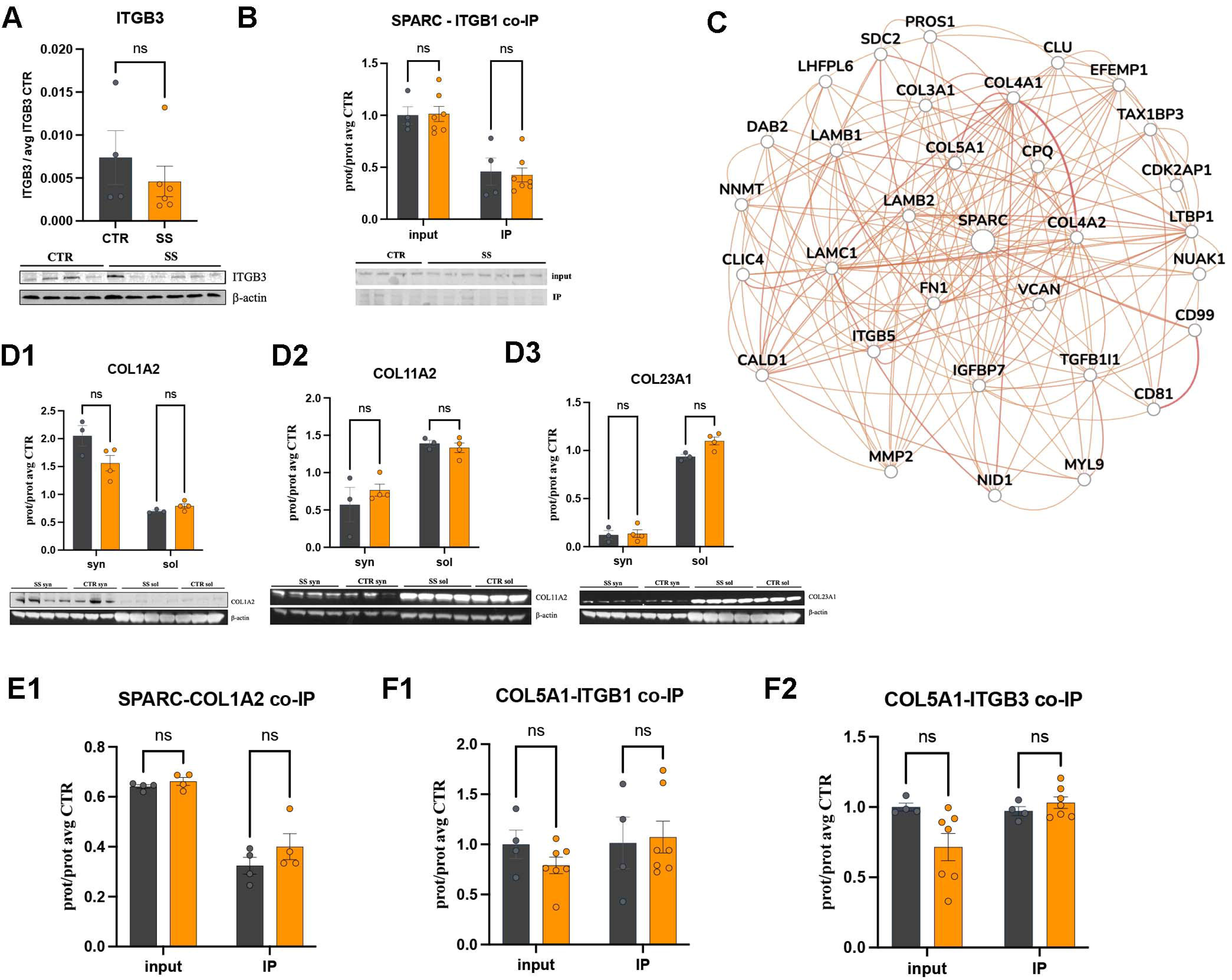
Protein-protein interactions are changed after social defeat stress. A) HumanBase SPARC interactome predicts interactions with multiple collagens (COL3A1, COL4A1, COL4A2, and COL5A1), fibronectin 1 (FN1), vitronectin (VTN), laminin 1 (LAMA1), versican (VCAN), integrin b5 (ITGB5) and others. B) WB analysis of the enrichment of ITGB1 shows no change between CTR and SS mice (p=0.3524). C) co-IP with SPARC and ITGB1 shows no change in interaction in SS mice compared to control (p=0.9931). D) WB analysis of the ECM protein enrichement in the synaptosomal and soluble fractions did not detect significant differential expression between CTR and SS mice, but differential enrichment of collagen 1A2 is in the synaptosomal fraction and COL11A2 and COL23A1 are significantly enriched in the soluble fraction. E-F) co-IP evaluation of the protein-protein interactions showed no changes for SPARC-COL1A2 (E) and COL5A1 with ITGB3 and ITGB1 (F).

**Supplemental Figure 4.**
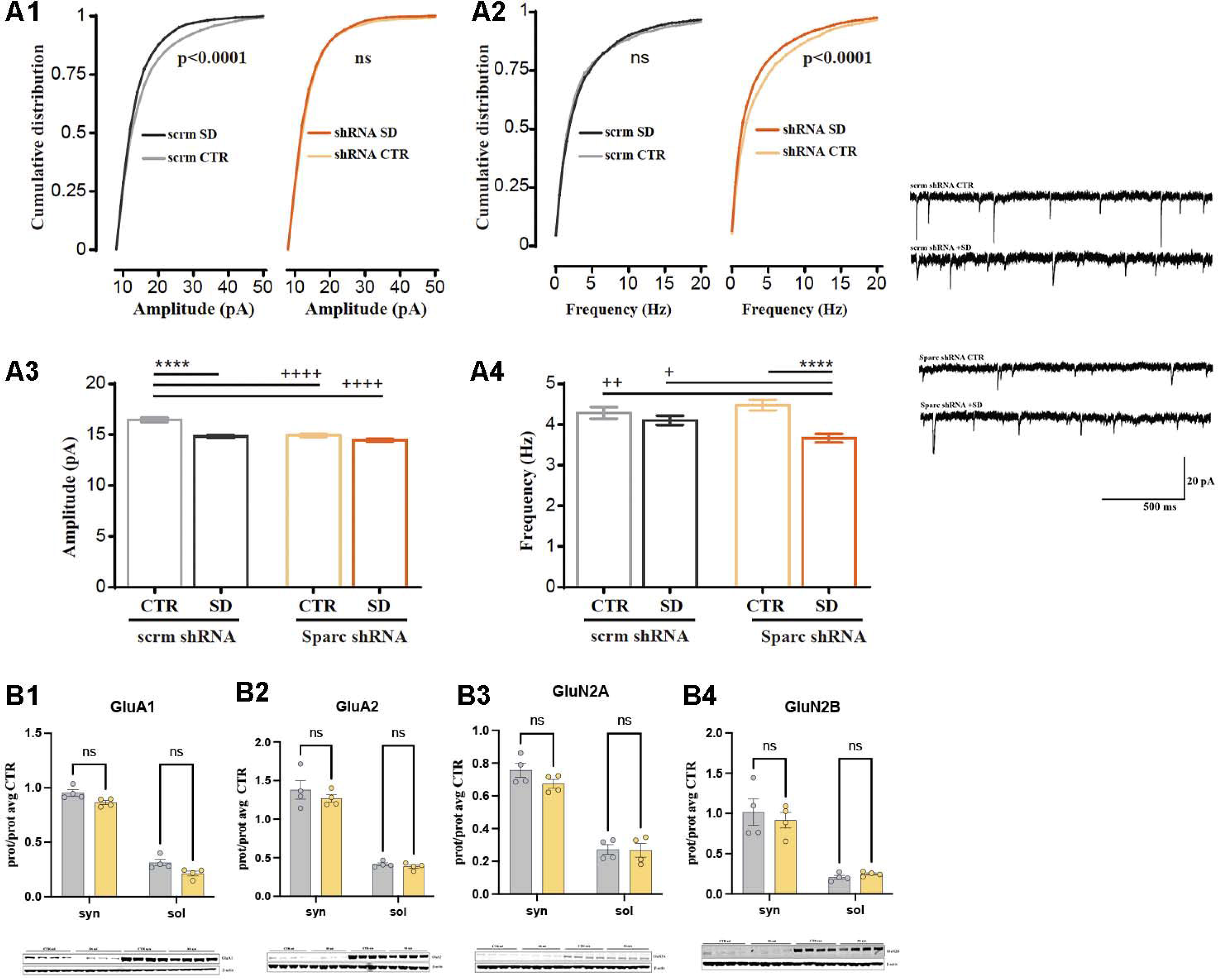
Synapse changes in mice with perturbed *Sparc* expression. A) Amplitude and frequency of excitatory postsynaptic currents are significantly decreased (p<0.0001) in WT mice with Sparc knockdown, but unchanged after social defeat. B) WB analysis of the glutamatergic receptors A1, A2, N2A, and N2B in synaptosomal fraction is unchanged in Sparc knockdown.

